# Physical cell-cell interactions regulate transcriptional programmes that control the responses of high grade serous ovarian cancer patients to therapy

**DOI:** 10.1101/2024.04.16.589574

**Authors:** Sodiq A. Hameed, Walter Kolch, Donal J. Brennan, Vadim Zhernovkov

**Affiliations:** Systems Biology Ireland, School of Medicine, University College Dublin, Belfield, Dublin 4, Ireland; Conway Institute of Biomolecular & Biomedical Research, University College Dublin, Belfield, Dublin 4, Ireland; UCD Gynaecological Oncology Group Catherine McAuley Research Centre Mater Misericordiae University Hospital, Eccles Street, Dublin 7

## Abstract

The tumour microenvironment is composed of a complex cellular network involving cancer, stromal and immune cells in dynamic interactions. A large proportion of this network relies on direct physical interactions between cells, which may impact patient responses to clinical therapy. Doublets in scRNA-seq are usually excluded from analysis. However, they may represent directly interacting cells. To decipher the physical interaction landscape in relation to clinical prognosis, we inferred a physical cell-cell interaction (PCI) network from ‘biological’ doublets in a scRNA-seq dataset of approximately 18,000 cells, obtained from 7 treatment-naive ovarian cancer patients. Focusing on cancer-stromal PCIs, we uncovered molecular interaction networks and transcriptional landscapes that stratified patients in respect to their clinical responses to standard therapy. Good responders featured PCIs involving immune cells interacting with other cell types including cancer cells. Poor responders lacked immune cell interactions, but showed a high enrichment of cancer-stromal PCIs. To explore the molecular differences between cancer-stromal PCIs between responders and non-responders, we identified correlating gene signatures. We constructed ligand-receptor interaction networks and identified associated downstream pathways. The reconstruction of gene regulatory networks and trajectory analysis revealed distinct transcription factor (TF) clusters and gene modules that separated doublet cells by clinical outcomes. Our results indicate (i) that transcriptional changes resulting from PCIs predict the response of ovarian cancer patients to standard therapy, (ii) that immune reactivity of the host against the tumour enhances the efficacy of therapy, and (iii) that cancer-stromal cell interaction can have a dual effect either supporting or inhibiting therapy responses.

## 1.0 INTRODUCTION

Ovarian cancer is the most lethal gynaecologic cancer causing over 180,000 deaths annually [1]. It represents a heterogeneous group of diseases which can be histologically divided into multiple subtypes [2]. Among these, high grade serous ovarian cancer (HGSOC) is the most common subtype, accounting for 80% of all ovarian cancer cases, affecting almost 240,000 women globally each year. Furthermore, HGSOC is the most lethal subtype, which is associated with poor long-term survival and acquired chemoresistance, linked to the extensive inter- and intratumoral heterogeneity as well as high chromosomal instability [3]. Also, HGSOC is molecularly characterised by TP53 mutations, frequent copy number aberrations, and defective homologous recombination DNA repair mechanisms [4]. Most patients present with advanced stage 3 or 4 cancer with a 5-year survival of approximately 30% [1]. Although HGSOC initially responds to platinum-based chemotherapy, acquired resistance means that recurrent disease becomes almost incurable, particularly in the platinum resistant setting where the median overall survival falls to approximately one year [4]. Therefore, it is imperative to understand the molecular dynamics underlying patient responses to therapy, in order to better stratify patients for effective therapy.

Cell-cell communication is fundamental in all body tissues and underlies diverse tissue structure, function, and processes in homeostasis, including immune responses. These interactions are dictated by complex biochemical signalling pathways, which regulate individual cell processes and intercellular relationships [5]. This communication can be perturbed in diseases leading to malfunctioning alongside dysregulated pathological scenarios [6]. Therefore, studies regarding cellular functions are increasingly considering the neighbourhood context of cells. Cell communications are mediated by a diverse range of molecules which include ions, metabolites, growth factors, hormones, integrins, receptors, structural proteins, junctional proteins, secreted proteins in the extracellular matrix. Indeed, some of these molecules (such as adhesion molecules) support structural interactions which bring about physical cell-cell interactions, while others (such as hormones, cytokines and growth factors) mediate cell communications over short or long distances [7].

Single cell RNA sequencing (scRNA-seq) allows an unbiased massive profiling of cells at single-cell resolution, thereby enabling diverse biological phenomena to be unravelled [8]. Indeed, many tools have been developed which harness scRNA-seq data for depicting cell-cell communication. These tools, which depend on prior knowledge-based curated ligand-receptor pairs, include CellPhoneDB [9], NicheNet [10], CellChat [11], SingleCellSignalR [12], and iTalk [13]. However, since scRNA-seq fails to conserve tissue architecture, the spatial context of cellular networks is lost in the process. Hence, these tools give no account of direct physical cell-cell interactions. Interestingly, other methods have been developed that can preserve physical cell-cell interactions. These are based on partial tissue dissociation to allow capturing and subsequent sequencing of paired cell multiplets, followed by multiplet deconvolution to identify the specific interacting cells. These include methods such as Clump-seq [14], CIM-seq [15], PIC-seq [16], pcRNA-seq [17] and ProximID [18]. Although these methods are efficient in deciphering physical cell interactions, they are technically demanding and require specialised personnel and equipment.

Doublets occur naturally at different rates in scRNA-seq when two cells are captured in a single reaction droplet. These occur largely due to incomplete tissue dissociation, therefore representing undissociated cells which are physically interacting with each other and end up being captured together as biological doublets. Of course, a proportion of doublets may also have occurred by random co-encapsulation of cells during scRNA-seq library preparation as technical artefacts. A recently developed method called Neighbor-seq which can be used to successfully identify and deconvolute the cellular composition of scRNA-seq ‘biological’ doublets in order to infer direct physical cell-cell interactions [19]. Here, we decipher the physical cell-to-cell interaction network in the HGSOC microenvironment using scRNA-seq doublets as the starting point. Focusing on cancer-stromal cell doublets, we unravel the communication mechanisms, explore the molecular dynamics underlying these interactions and decipher how these influence the patients’ clinical response to chemotherapy and cytoreductive surgery.

## 2.0 METHODOLOGY

### 2.1 Single-cell RNA sequencing data analysis

A pre-processed scRNA-seq count dataset and clinical information were obtained from a study by Olbrecht et al. [3], composed of samples from 7 treatment-naïve HGSOC patients, all subsequently undergoing platinum-based chemotherapy and four of them also cytoreductive surgery, collected from the ovaries, peritoneum and/or omentum. These patients were categorised into 3 clinical groups according to their response to first-line chemotherapy; complete remission (CR), partial remission (PR), and progressive disease (PD). Notably, 4 patients belong to the CR group, 2 patients belong to the PR group, and only 1 patient belongs to the PD group. The obtained count matrix of 20,483 cells was then processed using the Seurat package v4.3.0 in R v4.2.2, including only genes that occurred in a minimum of 10 cells and only cells that contained a minimum of 200 genes. After computing the percentage of mitochondrial gene content, cells which had a minimum of 300 and maximum of 5,000 total RNA counts as well as mitochondrial genes below 25% were considered for further analysis. The gene counts were log-normalised for the total expression per cell, scaled to a factor of 10,000, and the top 5,000 highly variable genes were selected for further analysis. A principal component analysis (PCA) of the top 5,000 highly variable genes was performed on the scaled data, and the first 20 principal components (PCs) were selected following an elbow plot of the top 40 PCAs. Next, cells were clustered using a graph-based nearest neighbour clustering algorithm at a resolution of 0.35 which has been reported to result in better identification of expected cell types [3]. The clusters were projected and visualised on a uniform manifold approximation and projection (UMAP) plot of the first 20 PCs. Cell type annotations were performed using a combination of reference-based and manual annotation methods. Reference based annotation was performed using the SingleR and Sctype R packages, followed by manual tuning to refine these automatically generated annotations based on the expression of canonical markers. Cell proportions based on tissues and clinical outcomes were computed and visualised using the ggplot2 R package.

### 2.2 Doublet deconvolution and modelling

Doublet deconvolution was performed using the Neighbor-seq deconvolution method, which is publicly available as an R package, with the default parameters. Essentially, the count matrix with the corresponding cell type annotation was provided as input. Through a series of steps, each barcode that represents a single reaction droplet in the dataset was modelled to predict the number of cells and the cellular composition using Neighbor-seq. Briefly, the algorithm first clusters together all cell types with the same annotation and prepares a marker gene set for each cell class using the top 100 genes. Next, synthetic doublets of all possible homotypic and heterotypic combinations of singlets are generated and a gene matrix is prepared by summing the gene sets of the constituent singlets. A random forest model is trained on an equal proportion of the singlets and synthetic doublets. Finally, the random forest model is then used to predict each barcode in the original dataset as either a singlet or doublet as well as the cellular composition based on probability scores [19].

Topic modelling was performed using the Latent Dirichlet Allocation (LDA) algorithm to explore gene expression changes occurring due to physical interactions in doublets using Python v3.9.12, following the steps developed by Pancheva et al. [6]. To model the cancer-fibroblast population an input matrix containing the singlet and doublet gene counts was used for a perplexity test to choose the appropriate number of k, setting a value of 2 to 20. Then, the LDA model was fitted first, for the singlet population to generate gene-topic and topic-cell probability matrices for k number of topics. A second LDA model was fitted for the doublet population using an additional 20 topics to generate a new probability matrix. The genes were then ranked based on the probability and the number of cells in each topic.

### 2.3 Survival analysis

Survival analyses were performed using the Survminer and Survival R packages. Ovarian cancer transcriptomic and clinical data were obtained by querying the Genomic Data Commons (GDC) bioportal (https://portal.gdc.cancer.gov/) using the RTCGA R Package [20] with a set of filtering parameters (project = ‘TCGA-OV’, data.category = ‘Transcriptome Profiling’, experimental.strategy = ‘RNA-Seq’, workflow.type = ‘STAR - Counts’, access = ‘open’). Data was further processed using the DESeq2 R package, and Cox Proportional Hazard regression model was fitted using the Survival package for all genes of interest. The genes, whose hazard ratio had a p-value < 0.05, were considered significantly associated with events/hazards. Next, a Log Rank test was performed for each of the significant genes (obtained from Cox regression output) to assess the clinical significance of individual genes using the survdiff function of the Survival package. To this end, gene expression cutoffs were estimated using the surv_cutpoint function based on maximally selected rank statistics, and categorised (as high/low) using the surv_categorize function of the Survminer package. Significant genes which had p-values < 0.05 from Log Rank test were visualised in Kaplan-Meier (KM) plots. The KM plots were fitted using the survfit function of the Survival package and visualised using the ggsurvplot function of the Survminer package.

### 2.4 Differential gene expression and Functional annotation

Differential expression of genes (DEG) was performed using the FindMarkers function of the Seurat package, and genes were considered significantly differentially expressed if the adjusted p-value (p-adj) < 0.05. Over-representation analyses were performed using the enrichR R package while gene set enrichment analyses (GSEA) were performed using the clusterProfiler R package. Enriched pathways were filtered for adjusted p-values < 0.05.

### 2.5 Ligand-receptor interaction analysis

Ligand-receptor analysis was performed using two methods:

1) Manual interaction scoring and filtering, the same method utilised by Halpern et al. [17] and Manco et al. [14]. To uncover ligand-receptor co-expression in cancer-stromal cell doublets, for each clinical outcome (CR, PR, PD), an average expression of each gene was calculated from the normalised expression data. A list of ligand-receptor pairs was obtained from Ramilowski et al. containing 708 unique ligands and 691 unique receptors [21]. An enrichment score 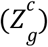 for each ligand or receptor gene (g) in a doublet cluster (c) was calculated using the formula:

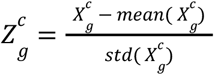

Where mean 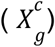 and std 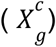 represent the overall mean and standard deviation across all 3 doublet clusters (CR, PR and PD). For each ligand-receptor pair (LRP) in a doublet cluster, an interaction score was computed as follows:

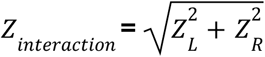

Where Z_L_ and Z_R_ represent ligand and receptor enrichment scores respectively. The obtained interactions were filtered through a set of parameters (average ligand gene expression > 0, average receptor gene expression > 0, interaction score > 1.5, ligand expressed in at least 5% of the doublet cells, and receptor expressed in at least 5% of doublet cells), and the final interactions were visualised in a network using Cytoscape v3.10.0.

2) BulkSignalR R package was also used to depict cell-cell interactions using a combination of ligand-receptor-pathway (LRPw) and the downstream signalling pathways/target genes, taking into consideration not only the expression of ligands and receptors but also the expression of the targets. Briefly, the dataset is decomposed into a triple of LRPw), in which for each LRP the expression of the downstream pathway target genes must be correlated and statistically significant to consider a LRP active. This tool scans the ligand-receptor database for information about LRPs, queries the KEGG, Gene Ontology Biological Process and Reactome pathways to delineate downstream pathways. Target genes are inferred from the reference network topology as those genes reachable from a receptor in a pathway by the directional edge annotated as “controls the expression of” [22]. LRPs were considered significant if the q-values were less than or equal to 0.01.

### 2.6 Transcriptional factor activity inference

The activities of transcription factors (TF) were inferred using a univariate linear model from the decoupleR R package [23]. The model allows the determination of active or inactive transcription factors based on the gene expression pattern in the dataset of interest. Essentially, this method achieves this by querying the collection of transcriptional regulatory interactions (CollecTRI) which consists of comprehensively curated TFs and their corresponding transcriptional targets obtained from 12 curated resources and available in the Omnipath database. So, for each sample/cell in the dataset and each transcriptional factor in the CollecTRI network (TF-genes), decoupleR fits a univariate linear model and scores each TF.

To allow for computational efficiency, the count matrix of the doublets were pseudobulked to sum up the gene expression of fibroblasts, cancer cells and cancer-stromal doublets at the sample level, using the aggregate.Matrix function of the Matrix.utils R package. The CollecTRI database list of TF-gene activity was manually obtained and a sample-level decoupleR was run with the default parameters. The top 25 significant TFs for each pseudo sample cluster were taken and then visualised in a heatmap across all clusters. Furthermore, to understand changes in TF activities across cell clusters, differential expression was performed at the individual cell level (without pseudobulking) as described in the previous section and decoupleR was run against the log2FC values of the significant DEGs, first for doublet vs doublets, and then doublet vs singlets. The top differentially active and inactive TFs were taken for each of the compared cell classes.

### 2.7 Trajectory inference

The scRNA-seq dataset was subsetted to obtain only doublet data. This was pre-processed through the Seurat scaling, clustering and UMAP projection using 10 PCs. Trajectory analysis was performed using the Monocle3 R package, which is based on the principle that cells in distinct biological states are characterised by functional differences in gene expression patterns that differentially define such states. This algorithm learns the overall gene expression trajectory in the scRNA-seq data and then places each cell in a position along the trajectory path based on its transcriptional state. Essentially, Monocle3 first reduces dimensionality and performs a projection using a UMAP method. Using the cluster_cells function, cells were clustered based on their gene profiles, and multiple partitions were built concurrently based on the connectivity of gene trajectories. Then, using the learn_graph function, cells were organised into trajectories based on reversed graph embedding and were projected onto the graph. To assign pseudotime values to each cell, the CR doublet cluster was chosen as the trajectory root and pseudotime values were obtained using the order_cells function. To identify genes that change along the pseudotime trajectory, a regression-based differential analysis was performed using the graph_test function and genes with a q-value <0.05 were filtered as significant genes. Genes with similar co-expression patterns were grouped into modules and associated with clinical outcomes using the find_gene_modules and aggregate_gene_expression functions. Functional annotation was performed for genes belonging to the modules associated with clinical outcomes, as explained in the previous sections.

## 3.0 RESULTS

### 3.1 scRNA-seq analysis identifies distinct cell types

A pre-processed scRNA-seq count matrix of 7 HGSOC patients from Olbrecht et al. [3] was analysed using the Seurat package. Following data filtering, normalisation, scaling and clustering, a total of 17,639 cells composed of 12,954 cells from tumour and 4,685 cells from normal tissues were retained. These cells were annotated first using reference-based automatic annotation tools, SingleR and Sctype R packages, followed by manual refinement based on canonical markers (figure 1a & supp Table 1). A total of 9 cell types were annotated, including macrophages, T cells (CD4+ and CD8+), dendritic cells (DC), natural killer cells (NK cells), B cells, Fibroblast/Stromal cells and cancer cells (figure 1b). As expected, normal tissues were dominated by fibroblast and stromal cells with a negligible cancer cell fraction (∼ 0.1%) while the tumour tissues had a large proportion of cancer cells and fibroblast/stromal cells, around 40% each (figure 1c, supp. Table 1). Interestingly, correlating cell proportion with clinical outcome revealed that only patients with complete remission had cells from normal tissues (figure 1d). Further, focusing only on the tumour tissues, patients with progressive disease (PD) had the highest proportion of cancer cells (∼80%) with the lowest immune cell fraction. Patients with complete remission (CR) had the lowest cancer cell proportion, most immune cells and highest fibroblast/stromal fractions. Generally, partial remission (PR) was intermediate in cell distributions, except for macrophages which were higher compared to both complete remission and progressive disease (Figure 1e, supp table 1).

**Figure 1:**
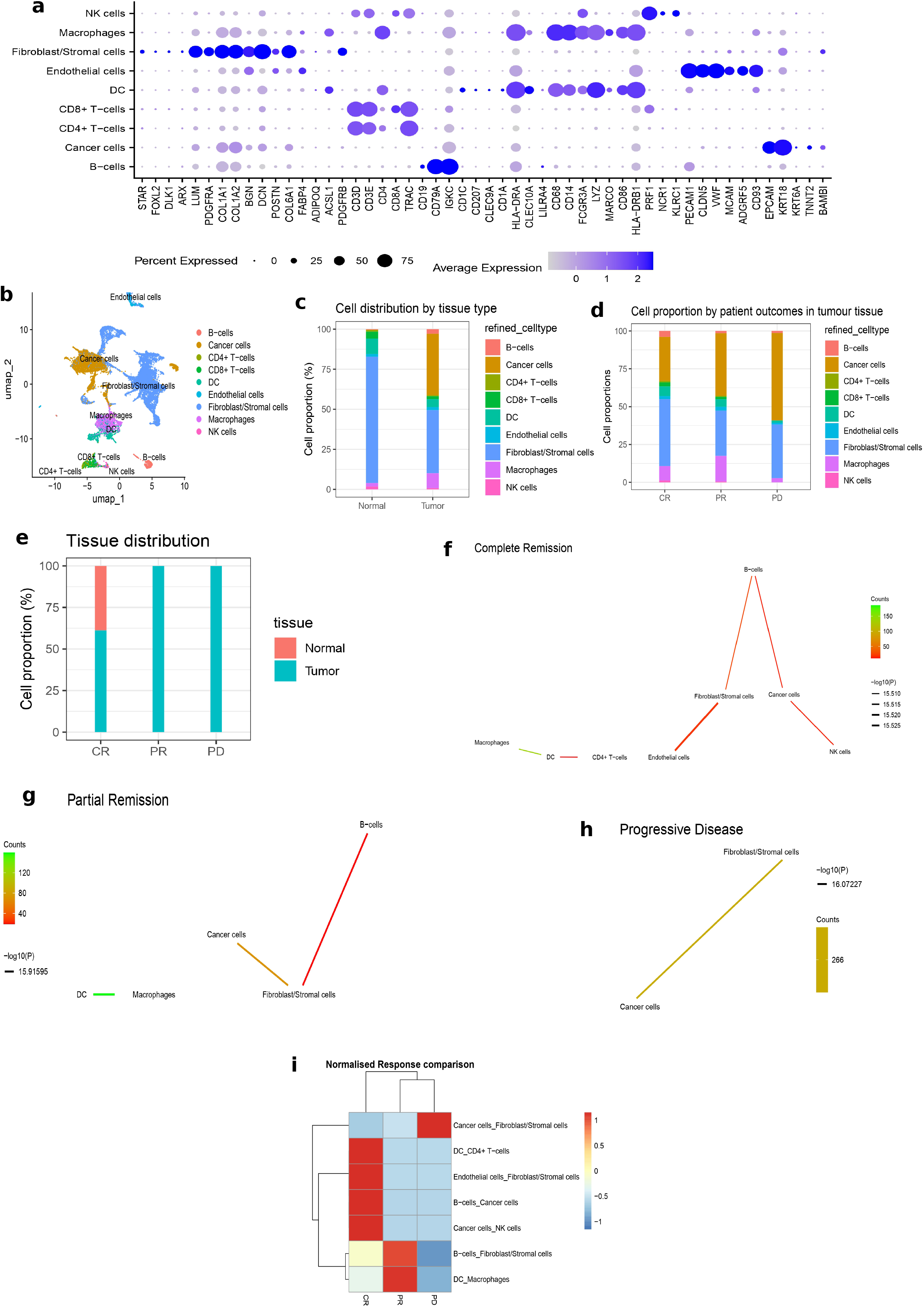
scRNA-seq cell annotation and doublet inference. a. Expression patterns of selected cell type-specific marker genes across cell types. b. Annotated cell types in the UMAP clusters (a) revealing 9 distinct cell types. Cell types were annotated using automatic annotation tools (SingleR and ScType) and manual refinement based on canonical cell type-specific markers (shown in 1a) c. Cell proportions in normal and tumour tissues. Normal tissues are dominated by fibroblasts and stromal cells, while tumour tissues contain more macrophages and a large number of tumour cells d. Proportion of normal and/or tumour tissues by clinical outcomes. Only complete remission samples contain cells from normal tissues. Partial remission and progressive disease samples consist of tumour tissues only. e. Cell distributions/proportions in tumour tissues from different clinical outcomes. The cancer cell fraction increases, while the immune cell fraction decreases as clinical outcomes worsen. f.-h. Physical cell interaction network obtained from doublets in patients with (f) progressive diseases, (g) complete remission, and (h) partial remission generated using the Neighbor-seq method. Lines are coloured by counts; red to green representing low to high counts respectively. i. Heatmap comparing physical cell interaction types by clinical outcome. The physical interaction (doublet) counts were normalised by the number of cells in each clinical outcome and scaled by a factor of 10,000. CR, complete remission; PD, progressive diseases; PR, partial remission

### 3.2 Doublet deconvolution identifies cancer-stromal physical interactions

The Neighbour-seq algorithm was utilised for doublet deconvolution. This assumes that doublets arise from physically interacting cells which fail to dissociate during single-cell library preparation and are captured in a single droplet as a result. This algorithm utilises a random forest model to predict the cellular composition of each barcode in a single-cell count matrix (see methods). The resulting doublet enrichment output was filtered to retain only heterotypic doublets (formed from 2 different cell types) with a minimum count of 10 and p-value <0.05, and then plotted in a cell-cell interaction network (figures 1f-h). The CR group had physical interactions involving immune cells such as cancer-NK cells, cancer-B cells, and macrophages-dendritic cells. There were also strong interactions between fibroblast-cancer cells (figure 1f). The PR group retained the macrophage-dendritic cell, fibroblast-cancer and B-cell-fibroblast interactions, but had no immune-cancer cell interactions (figure 1g).The PD group featured a significant enrichment of cancer-stromal interactions and no immune cell interactions present (figure 1h). A comparative overview of the interactions uncovered from the three therapeutic response groups is shown in figure 1i.

### 3.3 Cells in doublets communicate via ligand-receptor interactions with downstream signalling events

Ligand receptor co-expression analysis was performed as previously described. We initially used an interaction scoring approach to identify co-expressed LRPs, in doublets which may indicate that cell-cell communications occur. Overall, a total of 2165 potential LRPs were found for the CR, PR, and PD groups. The LRP list was filtered to retain only pairs with average ligand gene expression > 0, average receptor gene expression > 0, interaction score > 1.5, ligand expressed in at least 5% of the doublet cells, and receptor expressed in at least 5% of doublet cells. These thresholds retained only LRPs where both ligand and receptor were expressed in a reasonable proportion of cells. A total of 333, 69, and 188 LRPs were found for CR, PR and PD groups, respectively (supp table 1) and were visualised in Cytoscape (suppl figure 1).

Secondly, to understand the activeness of LRP pairs, BulkSignalR was used to model each doublet class to identify LRPs whose downstream pathways and target genes were also enriched. Essentially, this decomposes the dataset into triples of LRPw enrichments, whereby a triple is called as “active” when a statistically significant correlation exists in the expression of the ligand, receptor and the downstream target genes of the associated pathway (figure 2a). This resulted in the generation of LRPs and the associated downstream pathways (figure 2a, supp. figure 2a-b).. The CR and PR doublets displayed activities of LRPs whose downstream signalling events were involved in cell-cell interaction and immune-related events, among others. For example, the LRPs were those involved in cell-cell communication, cell junction organisation, and integrin cell-surface interactions, which emphasised ongoing interactions (figure 2a, supp. figure 2b). Also, immune-related events including antigen processing/presentation and MHC loading were enriched in the CR and PR group, an indication of a possible immune regulation of cancer progression. Paradoxically, the CR doublets contained CCL8/CCR5 and CCL8/CCR1 LRPs (supp. figure 2a), which induced downstream IL-10 signalling events (supp. figure 2b). CCL8 had previously been shown to be involved in the recruitment of T-regs and the promotion of tumour progression [24]. However, CCL8 in addition to CCL4/5 has been reported to induce the homing of CCR1/5+ intraepithelial CD8 T cells in ovarian cancer, thereby favouring improved response [25]. Similarly, the PR doublets interact through LRPs that induced downstream TGFβ signalling as well as IL-4/13 signalling events (figure 2a, supp. figure 2a-b); these have been reported to be associated with tumour promoting roles including growth, invasion and metastasis [26,27]. However, the PD doublets did not show an enrichment of LRPs for downstream junctional/interaction-related events, rather, there was an enrichment of “signalling by MET” which has been shown to promote invasive growth, survival and metastatic activities in cancer [28]. Also, similar to the PR doublets, there was an enrichment of LRPs that induced downstream IL-4/13 signalling (figure 2a, supp. figure 2a-b). Paradoxically, there was an enrichment of IL-15 signalling induced LRPs, which promotes anti-tumour immunity through NK-cell mediated tumour killing, although IL-15 has also been reported to have pro-tumoral activity in solid cancers [29]. Finally, we checked whether the LRPs obtained from the interaction scoring approach were present in the BulkSIgnalR outputs. This overlap identifies those LRPs which induce downstream target gene expression and hence can be deemed functional. Overall, 17, 16, and 5 LRPs of the previously scored LRPs were found for CR, PR, and PD doublets to overlap with the BulkSignalR outputs (supp figure 2c). Interestingly, these LRPs were those which drive many of the interesting downstream pathways discussed earlier (supp figure 2d).

**Figure 2:**
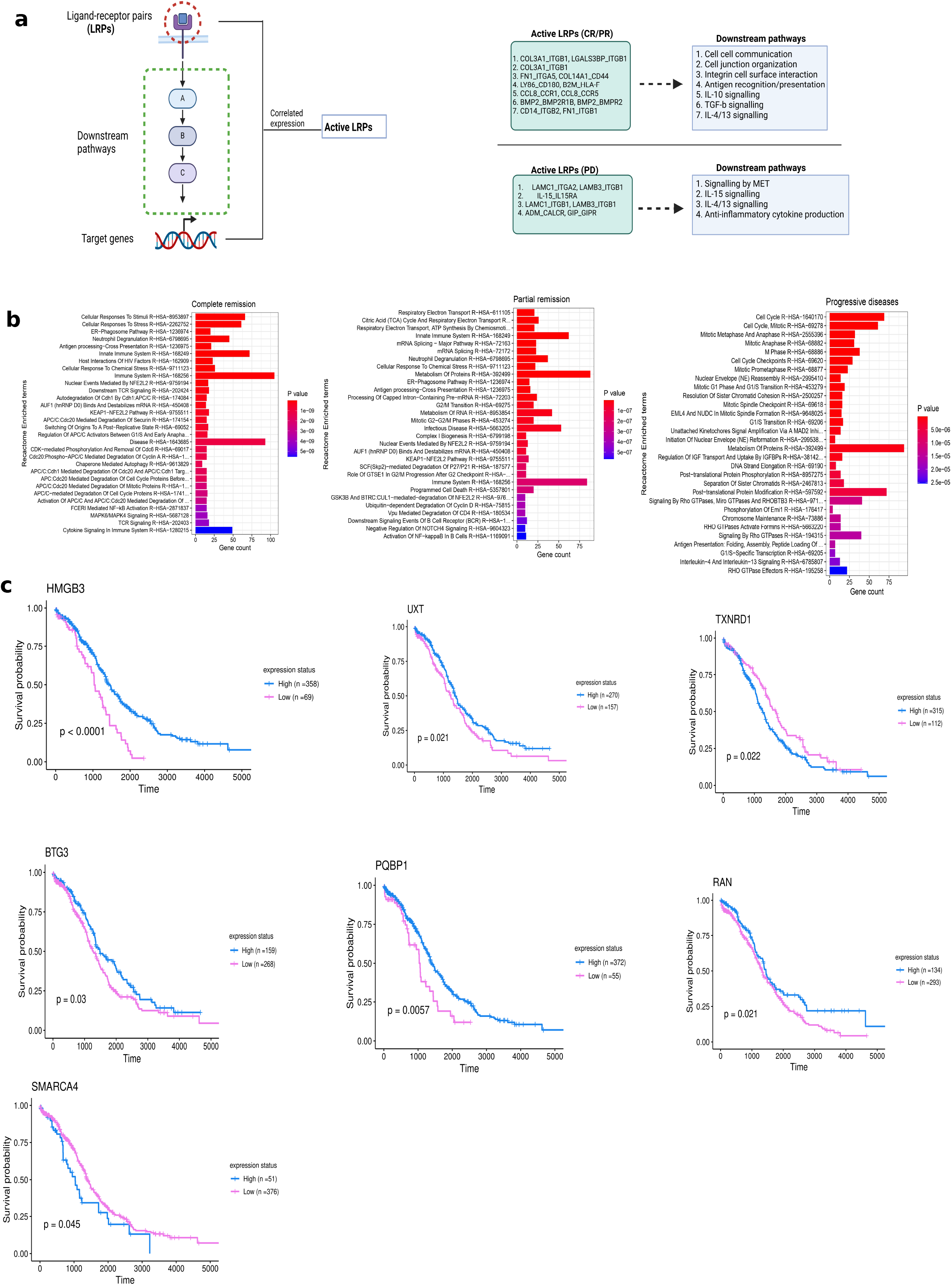
Cell-cell communication networks, functional annotation, and survival analysis. a. An overview of the BulkSignalR model workflow for predicting ligand-receptor interaction. For a ligand-receptor interaction pair to be considered active, the corresponding downstream signalling targets must have a significantly correlated expression level (left panel). Selected ‘active’ ligand-receptor pairs and the enriched downstream signalling pathways are displayed (right panel). CR, complete remission; PD, progressive diseases; PR, partial remission . b. Functional annotation of top-ranked LDA genes for complete remission (left), partial remission (middle), and progressive disease (right). Functional annotation was performed by querying the Reactome database for overrepresented pathways. Pathways were ordered by significance level (p-value), blue to red indicates lowest to highest p-values. c. Kaplan-Meier survival curves of ovarian cancer TCGA cohort for selected genes which were significantly associated with clinical prognosis from both Cox regression and Log Rank test. The bulk transcriptomic profiles of 429 ovarian cancer patients were utilised, for each gene of interest, patients were divided into low (purple line) and high expression (blue line) groups using the median expression value as the cut-off. Significance values obtained from the default log-rank tests are displayed in the plot as p-values

MP1_CD63

### 3.4 Topic modelling of doublets depicts interaction-related gene expression changes

Topic modelling using LDA was employed to model gene expression changes which occurred because of physical interactions between cancer and stromal cells, and to explain how this might have influenced clinical outcomes. LDA was fitted, first for the cancer and fibroblast/stromal cells singlets, and subsequently for the cancer-stroma doublet population. The genes were ranked based on the number of cells having the highest probability score for each doublet topic. For each topic, genes that occurred in less than 5 cells were removed and the top 30 genes with the highest number of cells were filtered for downstream analysis. A total of 564 genes for CR, 492 genes for PR, and 605 genes for PD were recovered for which expression has changed due to physical interactions (supp table 2). The topmost ranked genes uncovered for the PD doublets include CDCA3, PLK1, NEK2 and FAM64A (table 1) which are involved in ovarian cancer cell proliferation and drug resistance [30–33]. Furthermore, the topmost ranked genes in CR doublets included PRSS1, ECE1 and UBE2C, which promote ovarian cancer chemoresistance and progression [34–36]. Interestingly, MT1G, a tumour suppressor metalloprotein [37,38] was also upregulated in the CR doublets (table 1). Mechanistically, MT1G increases the stability of TP53 by inhibiting TP53 ubiquitination and increases the transcriptional activity of TP53 via direct interactions. To gain a broader overview of the overall upregulated genes, rather than individual gene functions, an over-representation analysis of functional annotation was performed on all LDA genes (figure 2b). This revealed that the CR genes recovered from LDA were genes enriched for cell cycle regulation-related (degradative) pathways, immune related processes involving antigen presentation/processing and cytokine signalling. The PD genes were highly enriched for proliferative pathways, RHO signalling, cell cycle, DNA replication and mitotic division, whereas the PR genes are enriched for both antigen processing/presentation and some level of cell cycle related pathways as well as pathways involving energy/protein metabolism.

**Table 1:**
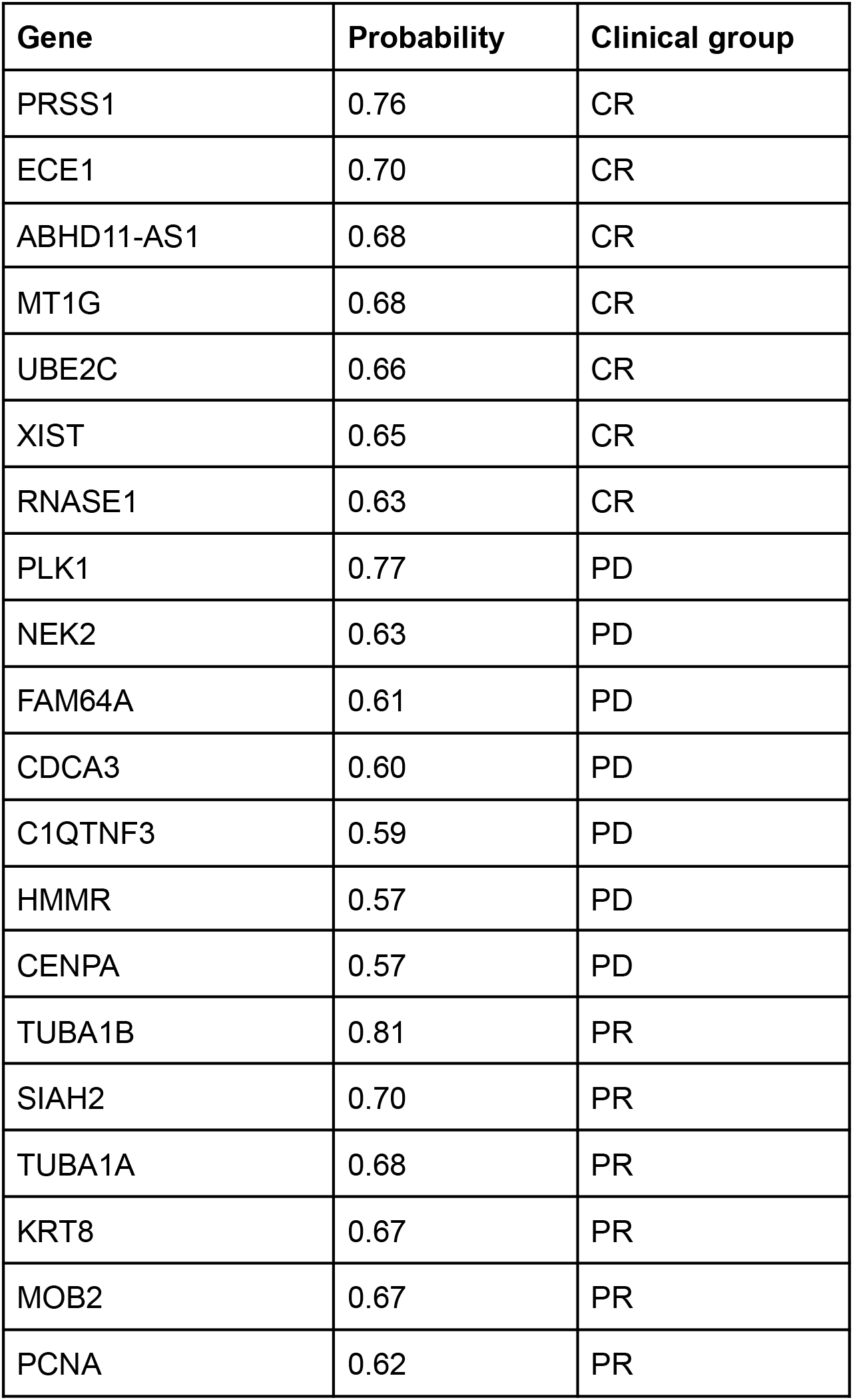
Top-ranked genes from Latent Dirichlet Allocation analysis. Latent Dirichlet Allocation was utilised to uncover genes that changed in expression due to interaction by segregation genes into latent unobserved topics. Probability scores represent the highest probability for each gene to have had a change in its expression in the doublet latent topic. CR, complete remission; PD, progressive diseases; PR, partial remission.

### 3.5 Doublet-specific LDA genes are associated with survival outcomes

#### 3.5 Doublet-specific Latent Dirichlet Allocation (LDA) genes are associated with survival outcomes

The LDA approach has not been widely utilised for cell-cell communication studies and thus required validation of its accuracy. LDA genes that are differentially expressed in the doublets compared to the singlets are likely to be uniquely associated with interactions. Therefore, a differential expression analysis was performed to identify genes uniquely expressed in doublets and to identify LDA genes that belong to these gene sets. For each clinical outcome (CR, PR, PD), the differentially expressed gene (DEG) analysis was performed on the cancer-stromal cell doublets first against the singlet population of stromal cells, and then against the singlet cancer cell population. Overlapping genes commonly upregulated compared to cancer and stromal singlets and part of the LDA genes, were thus identified as genes whose expression had increased due to cell-cell interactions. LDA does not describe the directionality of the changes in gene expression as to whether they were upregulated or downregulated, and although, the main objective of the original LDA model [6], was to depict upregulated genes, we reasoned that some of these genes may actually be downregulated compared to the singlets. Therefore, the DEG analysis was repeated to uncover LDA genes whose expressions were downregulated following interactions. Altogether, the differentially expressed LDA genes were combined as interaction-related genes. A total of 118 genes for the CR group, 116 genes for the PD group, and only 1 gene (KLK1) for the PR group were identified, which were considered as doublet-specific gene signatures (supp. table 2). To explore the clinical significance of these genes, a survival analysis was performed on all 235 genes using a TCGA cohort of 429 HGSOC cancer patients.

A Cox-regression analysis identified 55 genes whose expression was significantly associated with clinical outcomes (supp. table 2). Of these, 24 genes were associated with improved prognosis, while 31 were associated with poor prognosis, based on hazard ratios. Log Rank tests were performed on the 55 significant genes to assess the individual association of these genes on clinical prognosis. Here, 34 genes, including UBE2T, HMGB3, EMC9, ORC6, NEAT1, EIF3K, CRYAB, CCT2, GAPDH, MCM3, etc., were significantly associated with clinical prognosis (supp. table 2). KM curves produced for some of these genes (figure 2c) followed the same survival pattern as compared with the Cox hazard ratios.

These 34 genes were then visualised in the main dataset to explore how their expression truly changed in doublets as compared to singlets and in the 3 clinical outcomes. Indeed, of the genes associated with good prognosis, HMGB3, PQBP1, UXT, RAN, BTG3, and TXNRD1 were significantly higher in expression in the CR cancer-stromal doublets over their cancer cells and fibroblast singlet counterparts (supp figure 3). Although RAN and BTG3 were slightly upregulated in the PD doublets over their singlet counterparts, these were not as remarkable as those seen in the CR doublets. Taken together, these genes may be a probable contributing factor to the good patient outcome. Conversely, of the genes associated with poor prognosis, SMARCA4 was found to be significantly upregulated in the PD cancer-stromal doublets over their cancer and stromal singlet counterparts (supp figure 3). Also, NEAT1 (associated with poor prognosis) was significantly lower in expression in the CR doublets compared to their singlet counterparts (supp figure 3). Indeed, these findings are pointing to probable roles of SMARCA4 and NEAT1 in ovarian cancer disease progression.

**Figure 3:**
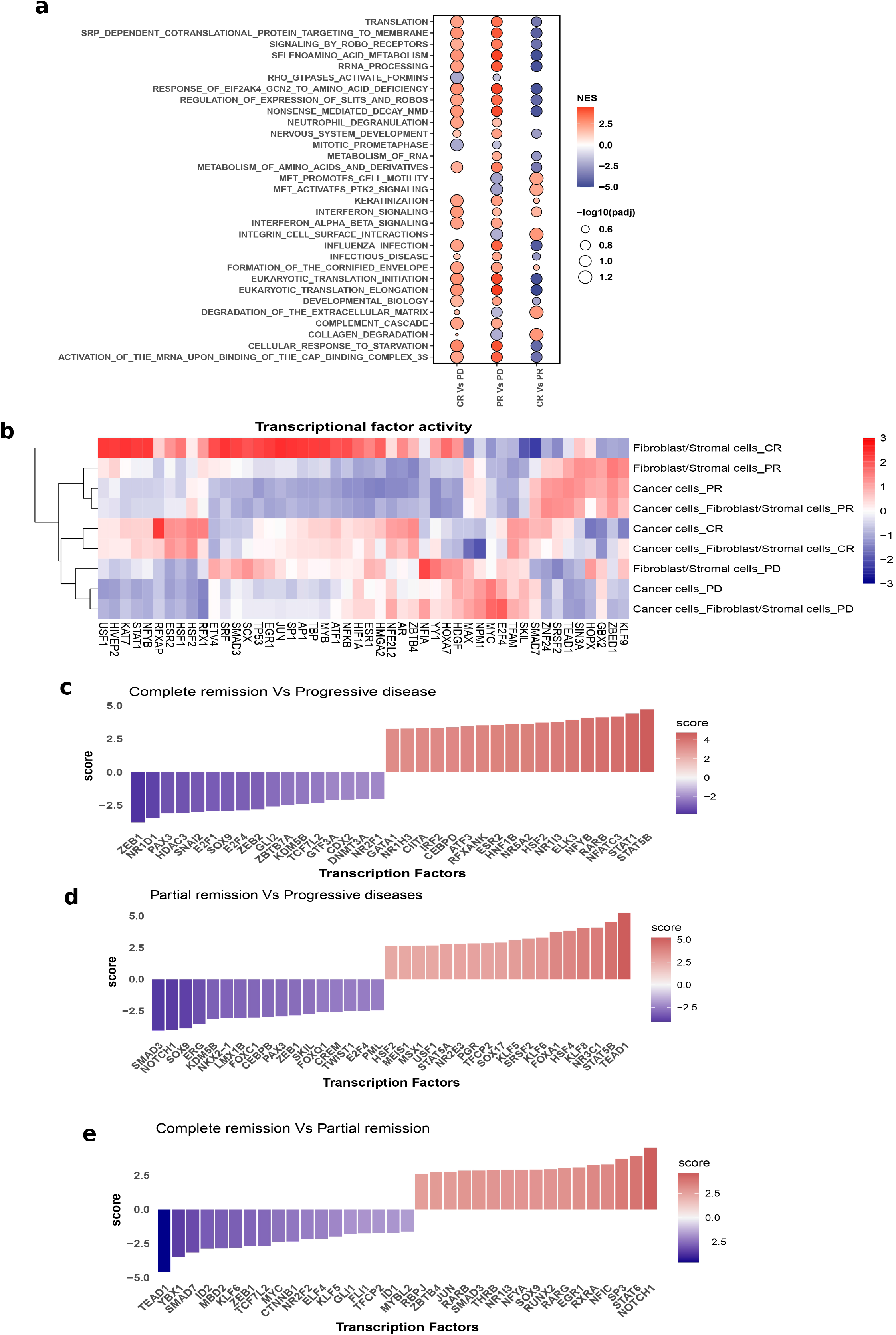
Functional annotation and transcription factor activity. a. Functional annotation of DEG when comparing doublets of different clinical outcomes b. Transcription factor (TF) activity across singlets and doublets showing clustering by clinical outcomes c-e. Differential TF activities in doublets for (c) complete remission compared to progressive disease (d) partial remission compared to progressive disease (e) complete remission compared to partial remission. Differential gene expression was initially performed for the compared cancer-stromal doublet groups and TF activities were inferred from the significantly differentially expressed genes.

### 3.6 Doublet classes show functional differences in gene expression

In order to understand gene expression changes that might have contributed to therapeutic response, we performed a classical differential gene expression analysis between the doublets, i.e. CR vs PD, PR vs PD, and CR vs PR doublets, giving rise to 3016, 2486, and 1081 significantly differential genes respectively. The significant DEGs were then functionally annotated by performing a pathway-level GSEA (figure 3a). Compared to the PD doublets, the CR and PR doublets showed a significant enrichment of pathways related to translation, protein processing and interferon signalling, whereas the PD doublets showed a significant enrichment of pathways related to MET signalling and mitotic cell division. Furthermore, comparing CR against PR doublets showed that the CR group had a more significant enrichment of the pathways related to translation, and protein processing, a possible indication that these proteins were possibly reducing proliferation, and/or perhaps stimulating immune functions, considering that interferon signalling was also enriched. However, CR also had an enrichment of MET signalling pathways compared to the PR doublets.

### 3.7 Epithelial-mesenchymal-transition events in doublets contribute to survival outcomes

Gene regulatory networks were assessed by inferring the transcription factor (TF) activities and how they change with cell states and clinical outcomes. To this end, the decouplR package was utilised. This draws on a curated TF-gene network, called the collection of transcriptional regulatory interactions (CollecTRI), which covers known TFs and the genes regulated by them, either positively or negatively. Mechanistically, this works by fitting a univariate linear model of TFs over the genes present in a dataset, drawing from the existing TF-regulatory network present in CollecTRI. TF activity was initially depicted across pseudo-samples of singlets (cancer cells and fibroblasts) and doublets (cancer-stromal cells) across clinical outcomes (CR, PR, PD), a total of 9 pseudo samples. Each pseudo-sample represents the aggregate of gene expression for all cells of the same type to form a single bulk RNAseq type of sample. As shown in figure 3b, there was a distinct transcriptional pattern for each clinical outcome as they formed separate clusters. Also, the singlet cancer cells clustered more closely with the doublet cancer-stromal cells, and this pattern was consistent across the 3 disease response groups. Next, we explored differential TF activities based on DEGs. Firstly, differential gene expression analysis was performed between the doublets as in the previous section, and TF activity was inferred from the fold changes (log2FC) of the DEGs. Compared to PD doublets, the CR doublets showed an enrichment of CIITA and IRF3 (figure 3c) which are related to MHC class II pathways [39][36] and interferon signalling response [40,41], respectively. In addition, CR had an enrichment of STAT1, which together with STAT2 is involved in downstream type-1 interferon signalling and has tumour suppressive roles [42]. Also, compared to both CR and PR (figure 3c-d), the PD doublets were enriched for epithelial-mesenchymal transition (EMT) activating TFs such as SOX9, ZEB1, ZEB2, TWIST1, and SNAI2. These TFs trigger EMT programmes which induce loss of epithelial adhesion and transformation into motile mesenchymal phenotypes, which enhance cancer metastasis, invasion, progression, and drug resistance [43,44]. Conversely, compared to PD doublets (figure 3c-d), both CR and PR activated STAT5A and/or STAT5B which can have dual roles as tumour promoting or tumour suppressing TFs [42]. Interestingly, comparing CR with PR (figure 3e) showed that PR doublets had an elevated activity of the inhibitors of differentiation (ID) TFs, ID1 and ID2, over the CR doublets. Indeed, ID1/2 play a role in tumour progression by enhancing cancer stem cell renewal, metastatic dissemination and proliferation [45]. Furthermore, the PR compared to the CR doublets featured an increased activity of the EMT activating the ZEB1 TF as well as MYC, which is a proto-oncogene involved in many aspects of tumorigenesis including tumour growth, survival, metastasis, and metabolism [46]. Similarly, the differential TF activities between doublets and their counterpart singlets were explored for the 3 clinical outcomes to understand the dynamics in gene regulatory networks that are driven by cell-cell interaction events and how these might have contributed to clinical outcomes. Here, a conventional DEG was performed between the cancer-stromal doublets and the singleton cancer cells and stromal cells. Interestingly, compared to their singlets, both PD and PR doublets had increased activity of NF-kB related TFs such as RELA, NFKB1 and NFKB [47], as well as the hypoxia-induced factor (HIF1A), and MYC (supp. figure 4). This indicates enhanced proliferation, possible cancer progression and reduced response to therapy [47–49]. Likewise, the activity of the tumour suppressor gene TP53 [50,51] was reduced in the PR doublets (supp. figure 4).

### 3.8 Trajectory inference

As discussed earlier, gene regulatory network analysis showed that differential TF activity is correlated with different clinical outcomes. Particularly, the interaction between cancer and stromal cells seems to play an important role across different disease stages. To evaluate whether cancer-stromal cell doublets have dynamic transcriptional states which transition as disease progresses along clinical outcomes, we performed a trajectory analysis using the Monocle3 package. Firstly, the cancer-stromal doublets were selected from the whole scRNA-seq dataset, pre-processed (see methods) and re-clustered, giving rise to 7 clusters that correlated with clinical response (figure 4a). Trajectory analysis projects cells in 2-dimensional UMAP space placing cell clusters in partitions based on transitions in transcriptional profiles. Cells which are connected by a transition in gene expression profiles belong to the same partition. We found the doublet clusters of the 3 clinical response groups well connected. When rooted by the CR cluster, a trajectory path could be constructed which branched; two branches remaining in the CR cluster and one branch connecting to the PR cluster. Similarly, the PR trajectory split into three branches; two remaining in the PR cluster, while the third led into the PD cluster (figure 4b). These results suggest that dynamic transcriptional transitions enable cancer-stromal cell doublets to adopt different cell fates that correlate with clinical outcomes. To identify genes that define these expression patterns, we grouped significantly differentially expressed genes with co-varying expression patterns into ‘modules’, which could be associated with doublets of different clinical outcomes (figure 4c). Overall, we found a total of 3062 genes enriched in the CR modules. These were genes functionally involved in interferon-alpha/beta/gamma signalling, translation, cytokine signalling and immune system related functions (figure 4d). On the other end of the spectrum, a total of 5779 genes were enriched in the PD modules, and these were genes that are functionally involved in cell cycle, mitotic division, DNA replication and RNA polymerase activity (figure 4d).

**Figure 4:**
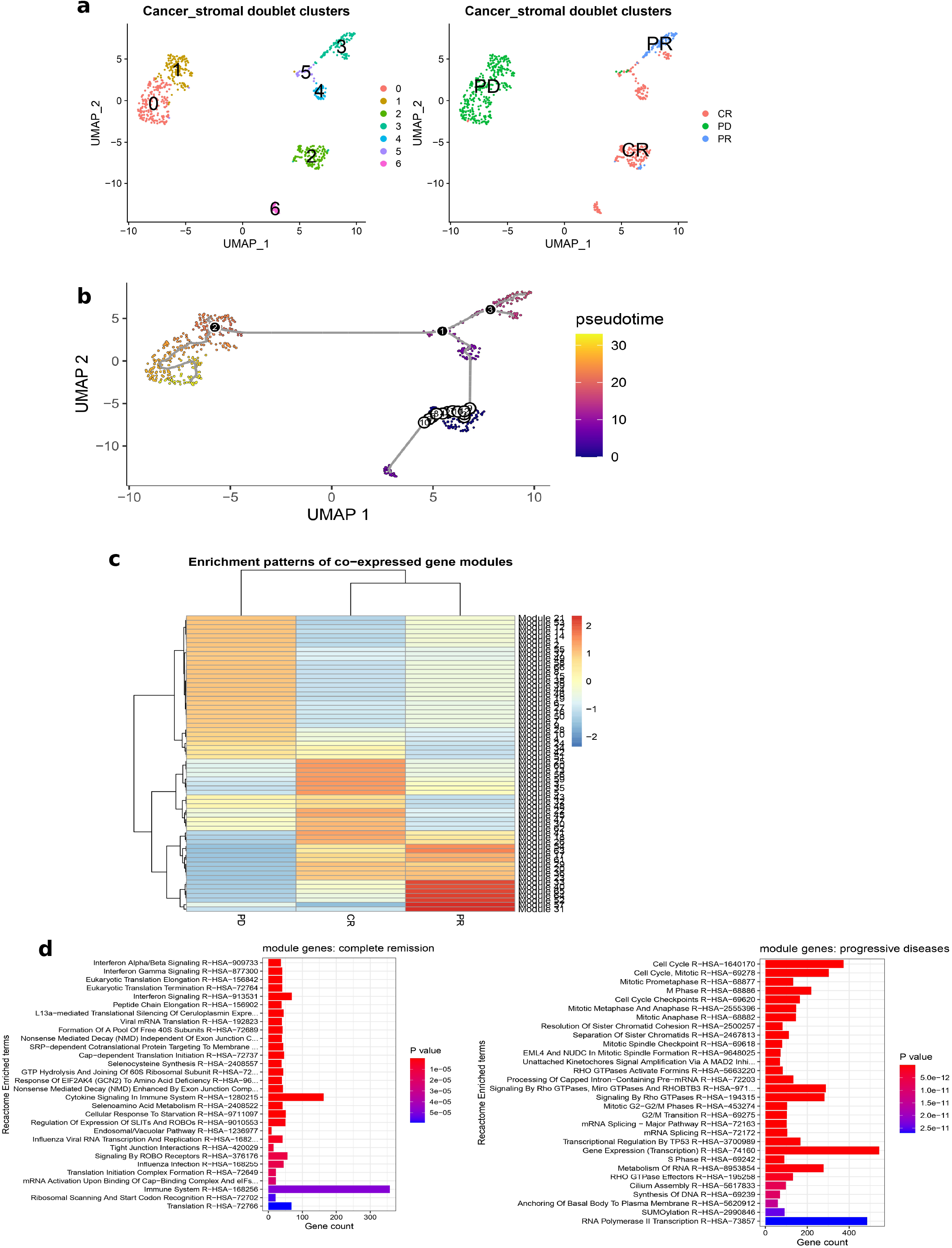
Trajectory analysis. a. Clustering of cancer-stromal doublets showing 7 distinct clusters (left panel) and cell clustering by clinical outcomes (right panel) b. Trajectory path across cell clusters. Circles represent the root cluster (starting point) of the path while black dots represent branching points. Cells are coloured along pseudotime which increases as cells become transcriptionally divergent from the root cluster such that cells closest to or at the root have low values and those furthest have the highest pseudotime values along the path. c. Heatmap of gene modules by doublet class. d. Functional annotation of the genes enriched in the CR modules (left) and the PD modules (right) CR; complete remission, PD; progressive disease, PR; partial remission

## 4.0 DISCUSSION

In most scRNA-seq studies, doublets are removed as technical artefacts that could confound data analysis. However, emerging studies suggest that a proportion of doublets could be cells which are physically interacting with each other and, therefore, fail to dissociate during scRNA-seq library preparation. Here, we focus on the physical interactions between cancer cells and stromal cells inferred from scRNA-seq doublets. Following cell annotation, we found the therapy responders (CR and PR), compared to non-responders (PD), had a good proportion of immune cells including CD8 and CD4 T cells, B cells and NK cells, especially in CR patients (figure 1e). This is indicative of immune infiltration, anti-cancer immunity and could have accounted for a better response to therapy. Indeed, tumour-infiltrating lymphocytes (TIL), particularly CD3+ CD8+ cells, have positive prognostic value in multiple cancers [52–55] including ovarian cancers [56], since they are involved in tumour antigen recognition and tumour killing. Indeed, the mere absence of immune cells in the PD cancers likely could allow tumour survival and cancer progression. Also, the microenvironment in the PD group was dominated by stromal cells, largely fibroblasts, which are known to influence cancer progression and invasion through direct or paracrine secretion of modulatory factors [57,58]. As expected, when physical cell-cell interactions were assessed from the scRNA-seq doublets, a good proportion of cancer-immune cell physical interactions were found in the responders, indicating an active anti-cancer immune response. In contrast, non-responders with progressive disease had no such interactions but were highly enriched in cancer-stromal cell interactions. This distribution was not surprising, considering the cellular profiles of the microenvironments uncovered earlier. These results confirm that the immune response to the tumour enhances the efficacy of chemotherapy and maybe also debulking surgery.

An interesting aspect was the modulatory effect of physically interacting cancer and stromal cells in relation to clinical outcomes, since these were present in both responders and non-responders. Indeed, these physically interacting cells communicate through a variety of mechanisms involving ligand-receptor pairs and downstream signalling pathways. Of interest, we found downstream signalling pathways specifically regulated by cell-cell interaction events, intriguingly including antigen processing/presentation related pathways downstream of ligand-receptor interactions (figure 2a). As cancer-stromal cell interactions are also strongly represented in responders, stromal cells may have a dual role. Besides their well-known role as immunosuppressors and promoters of cancer cell growth, there is also evidence that they can inhibit tumour growth by suppressing the immunosuppressive T regulatory cells in the tumour microenvironment [59]. Our results suggest that stromal cells in CR patients also may stimulate pathways in the cancer cells that contribute to tumour antigen presentation and enhanced cancer control by immune cells. However, considering that the tumour microenvironments of responders also had sufficient immune infiltration, coupled with the fact that there were physically interacting cancer-immune cell pairs, the enriched antigen processing/presentation may be a secondary effect of the immune milieu rather than a direct modulation by the stromal cells. For example, tumour-specific MHC-II expression on cancer cells is often induced in response to IFN-γ secreted by T cells in the tumour microenvironment [39]. Despite this, there were some LRPs whose downstream pathways involved IL-10, TGFβ, and/or IL-4/13 signalling across all doublets in responders and non-responders. Since these are typical immunoregulatory cytokine signalling pathways, it is probable that stromal cells have direct and differential effects on cancer cells in responders and non-responders. Gene expression regulated by physical cell-cell interactions included genes that are involved in cell cycle and mitotic division in non-responders, indicating that cell-cell interactions increased proliferative activity. Interestingly, the expression of cell cycle regulatory genes was also enriched in the CR group, but they also featured an enrichment of antigen processing/presentation related genes, suggesting that tumour cell proliferation is still taking place but is efficiently counteracted by immunosurveillance.

Interestingly, different transcriptional states of the doublets correlated with therapeutic response and survival GSEA revealed that doublets in responders were enriched in pathways of protein translation, Slit/Robo and Rho-Formin signalling (Figure 5a). Slits are ligands for Robo receptors which regulate neuronal guidance during development and in cancer can reduce adhesion and stimulate cell migration, but also suppress the formation of new blood vessels [60]. Formins are activated by Rho proteins and regulate the formation of cytoskeletal stress fibres strengthening focal adhesions and cell attachment [61]. This may indicate that maintaining adhesion and the integrity of the cytoskeleton contributes to the responder phenotype, and it is conceivable that this requires continuous protein translation. Additionally, responders showed an enrichment of interferon signalling, which aids anti-tumour immunity by stimulating immune cell function and also may be involved in controlling proliferation and progression in cancer cells [62–64]. Comparing the gene regulatory network of doublets in responders to non-responders showed that non-responders were enriched in the activity of core EMT activating TFs such as ZEB1/2, SNAI1/2, and TWIST1. Indeed, this further emphasises the potential drivers of the poor outcome as EMT events are associated with progression, metastasis and chemoresistance [40,41]. Contextually, this implies that stromal cells interact physically with cancer cells to induce EMT in the non-responders. Interestingly, a high expression of EMT genes has recently been linked to poor prognosis in ovarian cancer patients [65].

While our study makes some intriguing suggestions about the role of cell-cell interactions in the progression of ovarian cancer, it also has limitations. Firstly, the findings and conclusions are based on correlations, and were not validated experimentally. It would be worthwhile to experimentally validate salient in-silico findings in future studies, for example, using spatial transcriptomics on clinical samples to verify the type of cell-cell interactions, and 2D/3D co-culture models to study the molecular mechanisms in more detail. Secondly, due to the stringent dissociation protocol used, we only could analyse 700 doublets from 7 patients, however, the analysis was expanded using a larger TCGA cohort. Nonewisthanding, increasing the number of patients and doublets will increase the robustness and generalizability of the findings. Also, the genes and other changes found in the study had no clear directionality as to which cells specifically drove the changes observed, i.e. whether changes seen were cancer-specific or stroma-specific. Unfortunately, to the best of our knowledge there currently is no method that allows this distinction. Hence, the interpretations of some changes were heavily based on prior biological knowledge.

## 5.0 CONCLUSION

Overall, we have mapped changes in gene expression arising from binary physical interactions between cells in the tumour microenvironment. These changes correlate with responses to standard clinical therapy, and our analysis highlights some molecular mechanisms potentially contributing to the clinical outcomes. In particular, we discovered a dual role for cancer-stromal cell interactions, which were associated with tumour remission as well as with tumour progression. The gene expression states mediating such cell state changes could be mapped on a pseudo-time trajectory suggesting a progressive evolution of the non-responder from the responder phenotype. Taken together, our results provide an insight into the mechanisms involved in the modulation of cancer progression and/or evolution by stromal and immune cells through direct interactions with cancer cells.

## 6.0 ACKNOWLEDGEMENTS

This work was supported by Science Foundation Ireland (SFI) through grants 18/SPP/3522 and 22/FFP-A/10729. SH was supported through the SFI Research Training in Genomics Data Science, grant number 18/CRT/6214.

## SUPPLEMENTARY FIGURES

**supplementary figure 1: Ligand receptor interaction network**

A manual scoring approach was utilised to infer ligand receptor co-expression in

cancer-stromal doublets. Here, an average expression of each gene was calculated from the normalised expression data. A list of publicly available ligand-receptor pairs was obtained containing 708 unique ligands and 691 unique receptors (Ramilowski et al., 2015). An enrichment score for each ligand or receptor gene in a doublet cluster was calculated and filtered on certain parameters (see methods). The resulting ligand receptor pairs were visualised as a ligand-receptor interaction network for (a) complete remission, (b) partial remission, and (c) progressive diseases using Cytoscape v3.10.0.

**Supplementary figure 2: Inferred ligand-receptor pairs and corresponding downstream pathways**

a. Active ligand-receptor pairs (LRPs) inferred from cancer-stromal doublets for complete remission (left), partial remission (middle) and progressive disease (right) using BulkSignalR. For a ligand-receptor interaction pair to be considered active, the corresponding downstream signalling targets must have a significantly correlated expression level.

b. The downstream signalling pathways driven by the inferred LRPs (above) in the cancer-stromal doublets for complete remission (left), partial remission (middle) and progressive disease (right)

c. Filtered LRPs representing the overlap between the LRPs inferred above and those obtained using interaction scoring approach (see main text, section 3.3) for complete remission (left), partial remission (middle) and progressive disease (right).

d. The downstream signalling pathways driven by the filtered LRPs (above) in the cancer-stromal doublets for complete remission (left), partial remission (middle) and progressive disease (right).

**Supplementary figure 3: Expression pattern of genes in singlets and doublets of different clinical outcomes**.

Boxplots displaying the expression levels of some doublet-specific genes which were significantly associated with clinical prognosis from both Cox regression and Log Rank tests. A Kruskal-Wallis test was performed to compare expression across all groups while Wilcoxon tests were performed for pairwise comparisons between/within doublets and singlets. NS: non-significant; *: p < 0.05; **: p< 0.01; ***: p < 0.001.

**Supplementary figure 4: Transcription factor activity**

Differential TF activities of doublets compared to singlets for complete remission (top), partial remission (middle), and progressive disease (bottom). For each clinical outcome, a differential expression analysis was performed between the cancer-stromal doublets against the singlet cancer cells and stromal cells to obtain the differentially expressed genes. TF activities were then inferred from significantly differentially expressed genes. Significant genes are those with p-values < 0.05.

