## Supplementary material for "Physical cell-cell interactions regulate transcriptional programmes that control the responses of high grade serous ovarian cancer patients to therapy": Supllementary Figure 1

### SUPPLEMENTARY FIGURE 1

#### a. Complete remission (CR) network

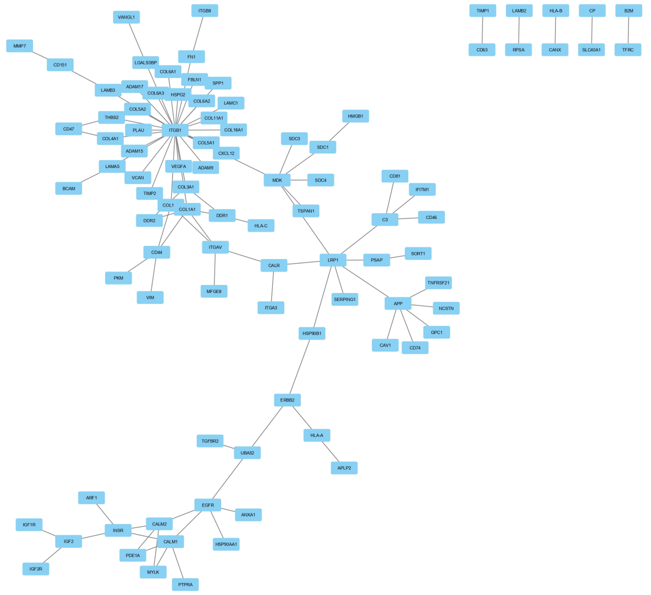


#### b. Partial remission (PR) network

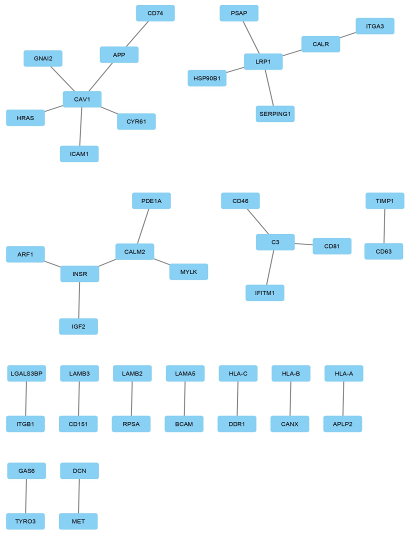


#### c. Progressive disease (PD) network

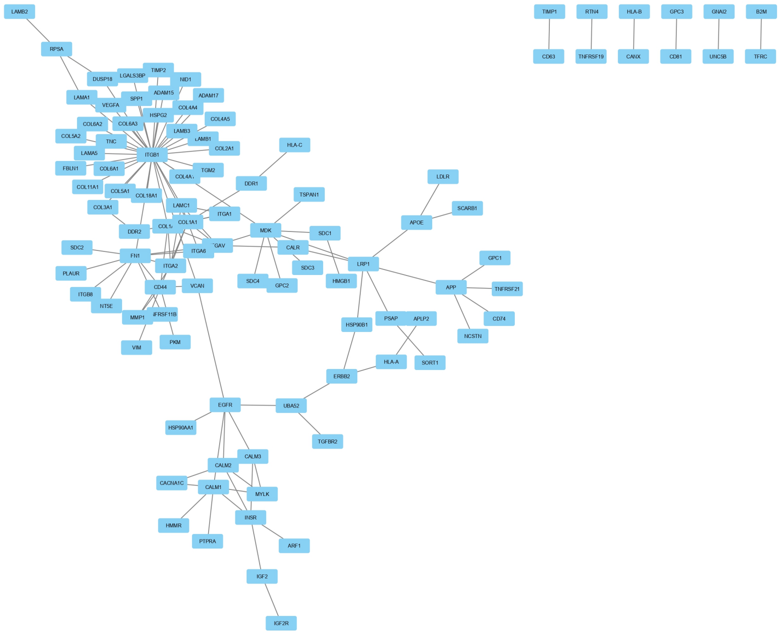
