## Supplementary figures and images for "Physical cell-cell interactions regulate transcriptional programmes that control the responses of high grade serous ovarian cancer patients to therapy"

### Supplementary Figure 2

#### SUPPLEMENTARY FIGURE 2

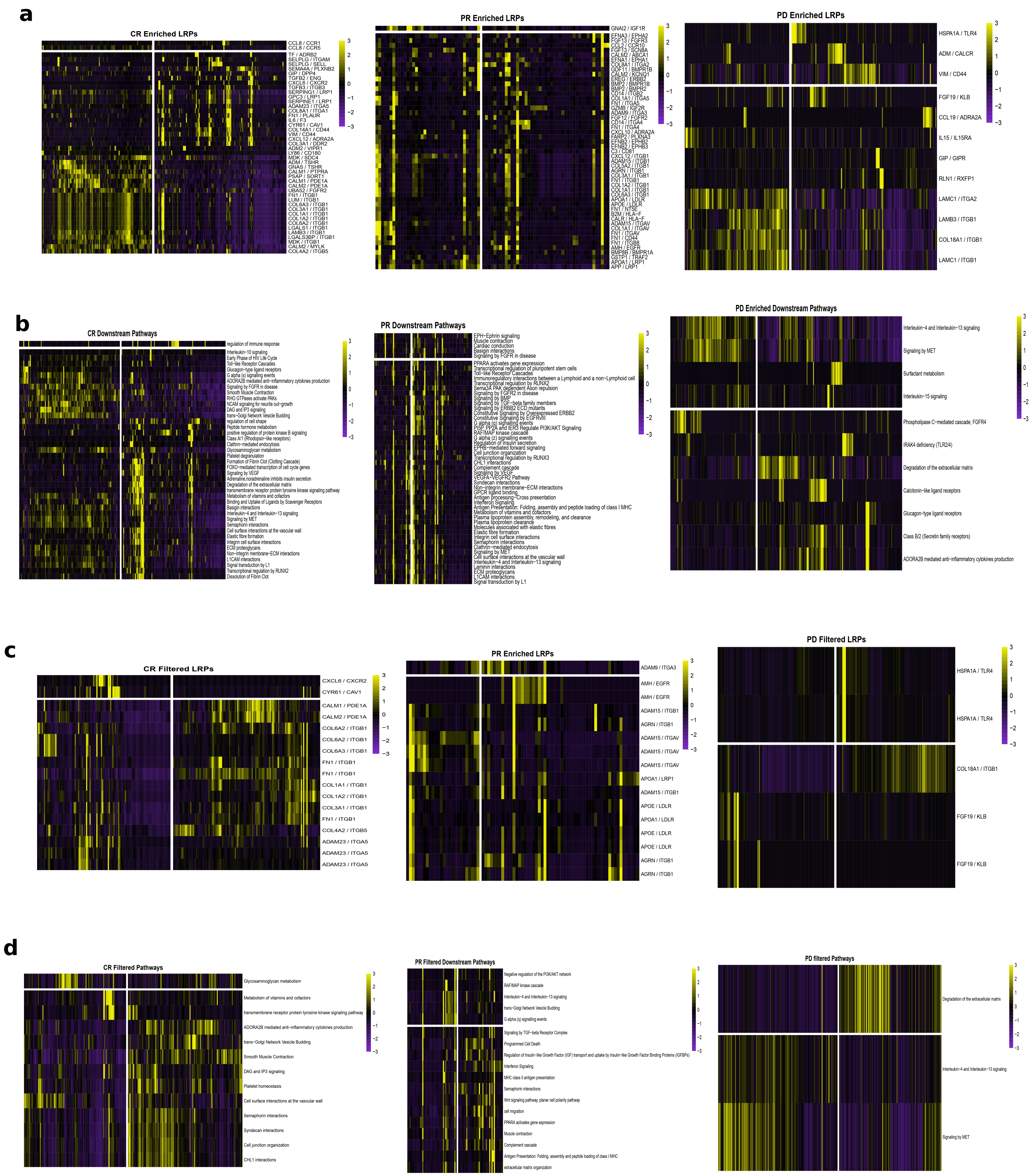

### Supplementary Figure 3

#### SUPPLEMENTARY FIGURE 3

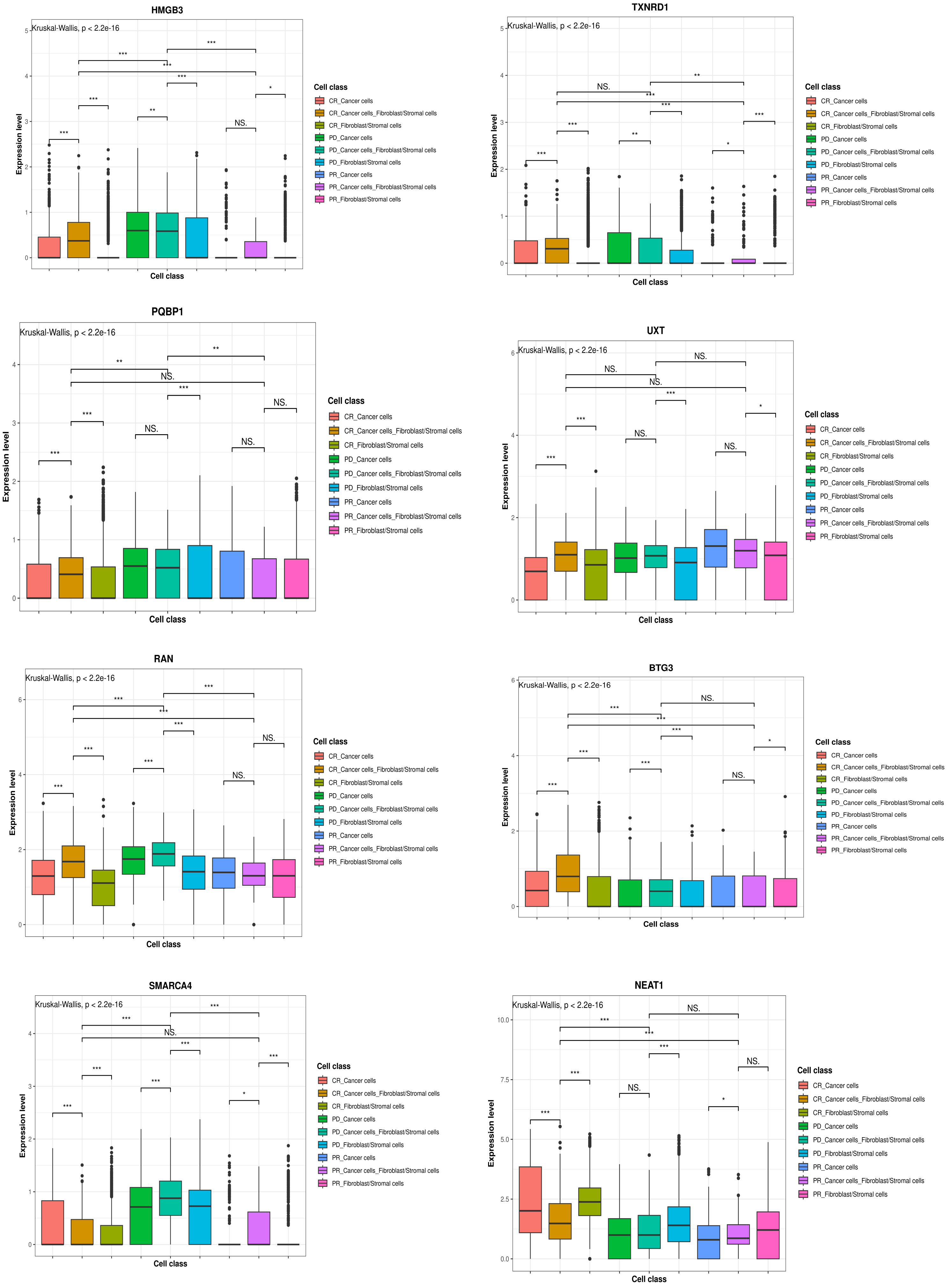

### Supplementary Figure 4

#### SUPLLEMENTARY FIGURE 4

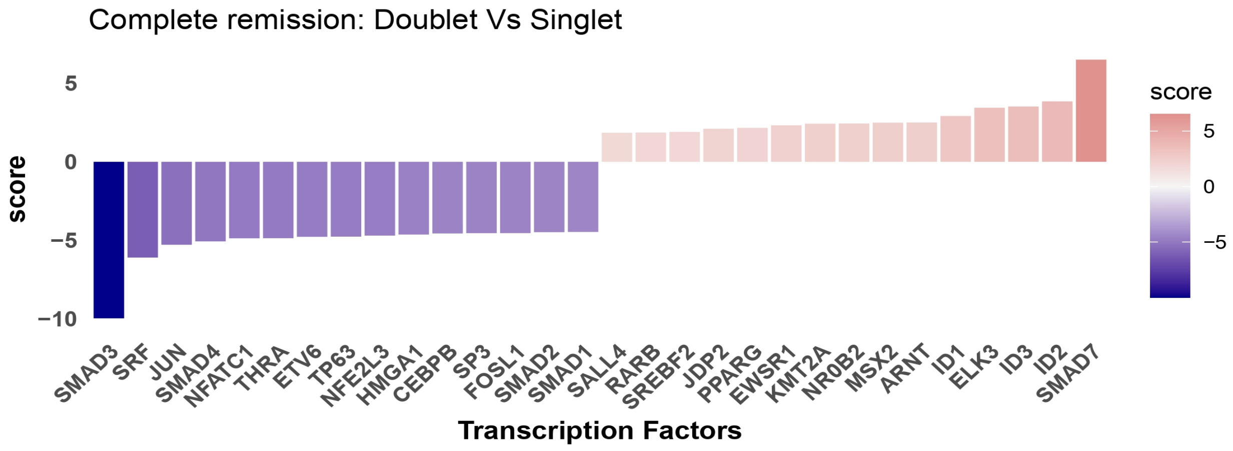


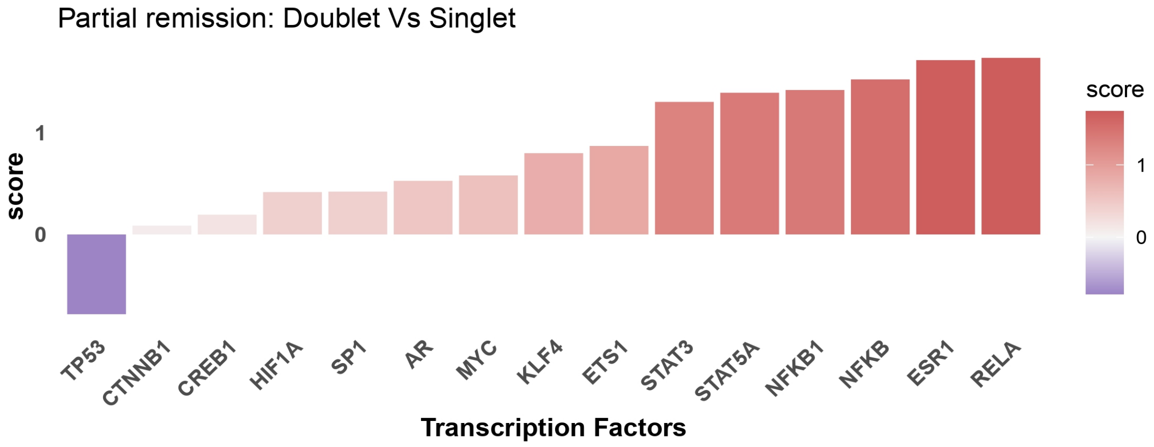


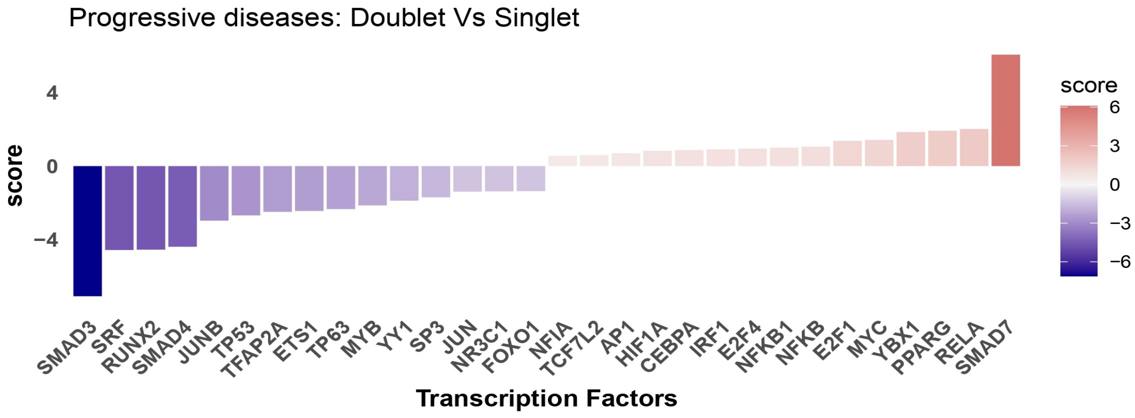
